# Characterizing Eastern spruce budworm’s large-scale dispersal events through flight behavior and stable isotope analyses

**DOI:** 10.1101/2022.10.20.513023

**Authors:** F. Dargent, J.N. Candau, K. Studens, K.H. Perrault, M.S. Reich, C.P. Bataille

## Abstract

Eastern spruce budworm moth (*Choristoneura fumiferana* (Clem.)) mass outbreaks have widespread economic and ecological consequences. A key explanation for the large-scale spread and synchronization of these outbreaks is the long-distance dispersal (up to 450km) of moths from hotspots (high-density populations) to lower-density areas. These events have proved difficult to monitor because dispersal flights occur only a few times a year, have no consistent routes, and commonly used tracking methods (e.g., population genetics, mark-recapture, radio telemetry) are inadequate for this system. Distinguishing between local and immigrant individuals is a crucial step in identifying the physical and ecological drivers of moth dispersal. Here we test whether isotopes of hydrogen (i.e., delta notation: *δ*^2^H) and strontium (i.e., strontium isotope ratios: ^87^Sr/^86^Sr), known to independently vary in space in a predictable manner, can be used to distinguish between local and immigrant adult spruce budworm moths. We used an automated pheromone trap system to collect individuals at six different sites in eastern Canada within and outside the current outbreak area of budworm moths. We first use moth flight behaviour and time of capture, currently the best available tool, to determine putative local vs. immigrant status, and then evaluate whether individual ^87^Sr/^86^Sr and *δ*^2^H differ between putative classes. At all sites, we detect immigrant individuals that differ significantly from putative locals. Saliently, sites where putative locals were sampled before the occurrence of potential immigration events (~10 days) showed the strongest differences between immigrant individuals and the locals ^87^Sr/^86^Sr and *δ*^2^H values. Sites where the collection of putative locals was close in time (hours) or following an immigration event had a less-clear distinction between putative immigrants and locals, and showed signs of mixing between these two groups. We speculate that recent immigration could have led to the misclassification of immigrants as putative locals. ^87^Sr/^86^Sr and *δ*^2^H data generally support the adequacy of current approaches using capture-time to detect immigration events, and provide enhanced resolution to distinguish between local and immigrant individuals. We discuss the broader implication of adding isotopes to the toolkit to monitor spruce budworm dispersal and suggest next steps in implementing these tools.

## Introduction

The eastern spruce budworm (*Choristoneura fumiferana* (Clem.)) is the most pervasive native pest of the North American boreal forest (MacLean, 2016). Large-scale outbreaks in eastern Canada occur in 30 to 40-year cycles (Jardon et al., 2003) and have severe ecological and socioeconomic consequences (Chang et al., 2012; MacLean, 2016). Spruce budworm (SBW) larvae preferentially feed on new foliage of balsam fir (*Abies balsamea* (Mill.)) and spruces (*Picea* sp.). Outbreaks can lead to high-density populations (>17×10^6^ large larvae per hectare) (Ludwig et al., 1978) which defoliate extensive areas of forest (e.g., approximately 52 million hectares in 1975) (Sleep et al., 2009). Multi-annual defoliation leads to high average tree mortalities (e.g. 85% in balsam fir stands), and growth reduction (e.g. of up to 90%), over extensive areas of forest (MacLean, 1980). In turn, this impacts forest regeneration and succession dynamics (Bouchard et al., 2007), increases CO_2_ emissions (Dymond et al., 2010; MacLean, 2019) and the likelihood of forest fires (James et al., 2017), and costs hundreds of millions of dollars in revenue loss and mitigation efforts (Natural Resources Canada, 2021).

Recent observations of the dynamics of rising budworm populations support a theory that attributed the rapid spread, extensive coverage, and widespread synchronization of spruce budworm outbreaks to dispersal events from high-density (i.e., outbreak) areas to low-density ones (Stedinger, 1984; Régnière et al., 2019a). At low-density sites, budworm population size is controlled by larvae mortality when failing to find food, and by predator and parasitoid-induced mortality, so that a limited number of adults contributes to the next generation (Stedinger, 1984). Immigrant arrivals can facilitate the transition to an outbreak density through the large influx of gravid moths causing a rapid population increase that in turn reduces stochasticity in larval survival caused during foliage searching, and cannot be suppressed by local enemies (Stedinger, 1984). Despite continuous research on this system for more than 50 years (Royama et al., 2017) and progress in control strategies (MacLean, 2019), gaps in our understanding of dispersal patterns and drivers make outbreaks a major challenge to predict and manage (Johns et al., 2019).

Understanding spruce budworm dispersal is challenging because dispersal events are irregular (Greenbank et al., 1980). Unlike classic roundtrip migrations or seasonal migrations that involve large numbers of insects following the same routes every year (e.g. Monarch butterflies (Urquhart and Urquhart, 1978;1979), Painted Ladies (Menchetti et al., 2019), Bogong moths (Warrant et al., 2016)), spruce budworm moths are mostly passive fliers and use wind currents for long-distance transport (Greenbank et al., 1980; Boulanger et al., 2017). After emergence, and for the ten- to fifteen-day life span of adult moths (Morris and Miller, 1954; Rhainds and Heard, 2015), whenever there is no rain and temperatures are above 14°C (Sanders et al., 1978; Greenbank et al., 1980), individuals will fly upward at dusk. These vertical flights can go up to an altitude of 400 m, with most individuals concentrated between 150 m and 300 m, and last for a few hours, after which the moths settle below canopy level for the rest of the day (Greenbank et al., 1980). During vertical flying, dispersal events may occur when moths are carried away by wind currents, and in some instances, hundreds of thousands of individuals can be transported hundreds of kilometres away by this process (Greenbank et al., 1980). These wind-facilitated dispersal events are relatively infrequent, occurring a few times per source site each year, and are directional, which means that simple modelling approaches that assume an isomorphic, radiation outside of a source area are inadequate, and could simultaneously involve large extents of forest as source and arrival locations. Such processes, in turn, make it challenging to predict the source of take-off, the direction of travel, or the landing area of dispersal flights. Additionally, this complex dispersal limits the viability of tracking approaches such as tagging (note that this approach is useful for monitoring local movement, i.e., distances below 100m (e.g. Sanders, 1983)) while the small size of this insect makes radio telemetry unfeasible. Finally, the lack of genetic structure across Eastern Canada’s local ‘subpopulations’ (Lumley et al., 2020) makes genetic-based monitoring approaches unviable. Consequently, there is currently no efficient tool to monitor spruce budworm dispersal, hampering our understanding of processes and conditions driving mass take-off and landing and ultimately limiting our ability to predict and manage the spread of outbreaks (Johns et al., 2019).

To improve monitoring of spruce budworm dispersal, a network of 22-30 automated traps has been deployed by the Federal and Provincial governments of Canada (Canadian Forest Service and the provinces of Quebec, Nova Scotia, and Newfoundland and Labrador) across Eastern Canada since 2018. This network is strategically set up to sample populations within outbreak areas and at the margins of the SBW distribution, where density ought to be low but susceptible to the spread of the outbreak (Figure 1). To gather information that can be used to distinguish between local activity and immigration events, these traps lure SBW with pheromones, and once an individual enters the trap it gets stuck to a roll of sticky paper. At four times during the day, pictures of the individuals caught in the sticky paper are transmitted to an online platform and moths are recorded as either new or previously-present captures. The time frame of the captured photographs is used to inform whether individuals are to be interpreted as potential locals or immigrants. Crucially, local individuals are expected to fly around dusk (Greenbank et al., 1980), thus, captures occurring between 17:00 to 23:00 are interpreted as potential locals. Immigrant individuals are expected to have some delay between their local-flying time and the time at which they land in a non-local patch after being carried away by winds. Therefore, captures between 23:00 and 05:00 are interpreted as putative immigrants. To enhance the accuracy of this approach, other lines of evidence are considered when trying to determine the migration status of moths, these include, (1) local phenology, which informs if conditions are adequate for adults to be present in the area; (2) temperature and wind patterns at the time of capture, which informs whether moths are likely to have been flying at a given site and the potential trajectory of travel; and (3) radar data, which can detect when large numbers of moths disperse. Although this multifaceted approach improves the detection of dispersal events, it has some potentially important limitations. On the one hand, short-distance migrants could arrive a few hours after their vertical flight and before the 23:00 cut-off point, which would lead to mixing between presumed locals and short-distance immigrants. On the other hand, after an immigration event, previous-day immigrants would initiate their vertical flight behaviour around dusk, and therefore captures between 17:00 and 23:00 may be composed of a mixture of locals and immigrants. One way in which the accuracy of these methods can be validated, and lead to improved detection of immigration events, is by looking at the isotopic signals of captured individuals.

**Figure 1:**
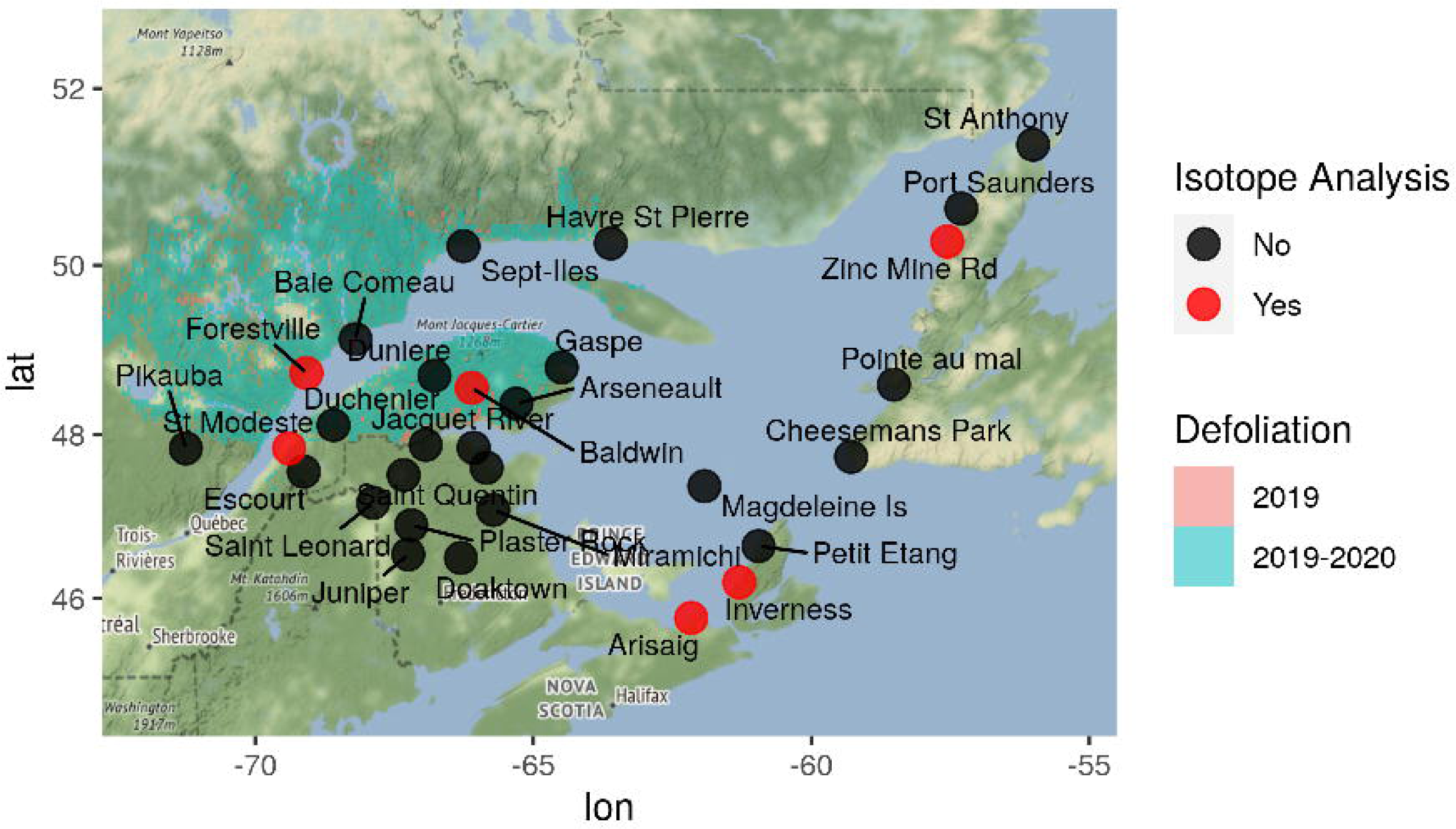
Trap location and eastern spruce budworm (SBW) defoliation area. Map of eastern Canada with location of automatic traps in the network (black points), traps used in this study (red points) and the 2019 and 2020 SBW area of defoliation, which is used as a proxy for population density and extent of the outbreak in a given year.

Isotopes of hydrogen (referred to using the delta notation or *δ*^2^H values) and strontium (referred to using the ratio of ^87^Sr to ^86^Sr or ^87^Sr/^86^Sr) are intrinsic markers of geographic origin because they reflect where an organism developed, and thus can identify whether an individual grew up in the area where it was captured (Wassenaar and Hobson, 1998; Hobson, 1999; Flockhart et al., 2015). Local isotopic signals are integrated into an organism through the process of tissue development. As an organism grows and forms new tissue, hydrogen and strontium isotopes are incorporated into its body from the local food and water they consumed (Bowen et al., 2005; Bataille et al., 2020). Studies on migratory insects (Flockhart et al., 2015Reich et al., this issue; Lindroos et al., this issue), demonstrate that tissues with low metabolic activity (e.g., wings) will carry and preserve the isotopic signal of the location at which SBW larvae fed on spruce or balsam fir needles (Lindroos et al., *this issue*). As SBW moths do not feed during their adult life and live a short adult life (i.e., 10-15 days), it is likely that wings and other tissues preserve the isotopic signature of the larval stages. Hydrogen isotopes are used in insect population ecology because of the predictable relationship between chitinous wings, which have low metabolic activity (Lindroos et al., *this issue*), and precipitation *δ*^2^H values (Hobson, 2019). The *δ*^2^H values in precipitation vary predictably in space and time with the hydrological cycle– in particular through the influence of evaporation and precipitation processes which in turn leads to the differential concentration of heavy relative to light hydrogen in local water (i.e., fractionation) (Bowen and Revenaugh, 2003; Bowen et al., 2005). *δ*^2^H values variation in organisms follows those of precipitation but also includes a predictable contribution of physical and metabolic processes that segregate light vs. heavy isotopes further (Bowen and West, 2019). The ^87^Sr/^86^Sr ratio is another tool with strong potential to complement hydrogen isotopes for ecological provenance studies (Bataille et al., 2020). ^87^Sr/^86^Sr ratios in ecosystems usually vary at higher spatial resolution than *δ*^2^H values following the chemical composition and age of surface geology, are stable through time and are corrected for metabolic-induced fractionation (Bataille et al., 2020). As such, ^87^Sr/^86^Sr ratios vary independently from *δ*^2^H values and have a strong potential to provide more specific geolocation potential (Bataille et al., 2020). Recent progress in analytical methods has allowed the successful application of ^87^Sr/^86^Sr ratios in wings to track the migration of other Lepidoptera (Reich et al., 2021). However, SBW are small insects and their wings are too small for a precise ^87^Sr/^86^Sr analysis. While Sr concentrations and ^87^Sr/^86^Sr ratios in the wings of migratory insects have been shown to reflect the larval diet and local soils, it is possible that some strontium might cycle through insect bodies (Reich et al., *this issue*). However, for SBW, adults don’t feed which means that there is no addition of dietary Sr during the adult stage. Additionally, SBW moths live for approximately 10 days following emergence (Greenbank et al., 1980) and up to 4 days after immigration (Rhainds et al., 2022) limiting the possibility of Sr intake from locations different from the larval stage site. Consequently, it is likely that the ^87^Sr/^86^Sr ratios in short-lived SBW reflect the larval stage and can be applied for geolocation.

Eastern Canada is an excellent location to apply insect isotope geolocation techniques. The climate of eastern Canada is varied with strong precipitation and temperature gradients going from the coast to more in-land locations leading to large *δ*^2^H value variations across the breeding range of eastern spruce budworm. Similarly, the geology of Eastern Canada is one of the most varied in the world with Paleozoic carbonate units juxtaposed with Precambrian cratons and Grenville orogeny lithologies (Canada and Wheeler, 1996) leading to large ^87^Sr/^86^Sr ratios variability across the breeding range of eastern spruce budworms. Consequently, these geolocation tools are highly applicable across this study area. Combining hydrogen and strontium isotopes have shown great promise in increasing the specificity of geographic assignments of animals (Reich et al., 2021). Dual hydrogen-strontium isotope geolocation has a high potential to facilitate the identification of local vs. immigrant insects (i.e., nominal assignment) as well as estimating the location of origin of immigrant individuals (i.e., continuous geographic assignment) (Wunder, 2010; Ma et al., 2020; Magozzi et al., 2021). As a first step towards modelling the dispersal dynamics of spruce budworm, we test the applicability of *δ*^2^H values and ^87^Sr/^86^Sr ratios as tools to distinguish between local and migrant individuals of eastern spruce budworm moths in six automated trap sites across the current outbreak and potential expansion distribution of the species (Figure 1). These automated trap sites are chosen based on the current spread of the spruce budworm outbreak. Three of our sites are in locations undergoing outbreak dynamics and thus have high population densities (Forestville, St. Modeste and Baldwin in Quebec, sampled in 2019), and the three other sites are in areas susceptible to the spread of an outbreak but remained at endemic levels with low-density populations at the time of sampling (Arisaig and Inverness in Nova Scotia 2020, and Zinc Mine in Newfoundland in 2019). First, we use a subset of individuals captured at these sites, presumed to be locals, and individuals following an immigration event that are presumed to be immigrants. We hypothesise that selected individuals will differ in their *δ*^2^H values and ^87^Sr/^86^Sr ratios given that locals and immigrants likely fed and formed their tissues in different localities of eastern Canada. Second, we discuss currently used flight-behaviour approaches to determine the origin of individuals, contrast them to isotopic approaches, and make suggestions for better understanding dispersal of SBW.

## Materials and Methods

### Determining local vs. immigrant individuals through flight behaviour, phenology and atmospheric modelling

Our automatic trap network uses automatic pheromone traps (Trapview+ model; EFOS d.o.o., Slovenia). Each trap has a synthetic pheromone lure that attracts male spruce budworm moths which then fall onto a roll of sticky paper. Four times a day at approximately 05:00, 11:00, 17:00, and 23:00 the trap cameras take a picture of the moths on the sticky roll and upload the image to a server via a cell phone network connection. The images are available in real-time and the sticky paper can be rolled remotely if it has a high number of moths attached. Moth numbers are automatically estimated by the Trapview software (EFOS d.o.o., Slovenia), and then validated by CFS personnel to confirm proper species identification. At the end of the flight season, the full roll is recovered and stored under laboratory conditions. A sticky roll can be later unrolled and the chronology of moths captured on the paper can be established so moths captured at a specified date and time can be recovered for isotope analysis (Supplementary Figure 1).

Moths are presumed as local or immigrant based on their flight behavior and the time of capture. Individuals captured between 17:00 and 23:00 are assumed to be locals, because SBW moths tend to initiate their flight activity around dusk (Greenbank et al., 1980; Régnière et al., 2019b). Moths captured between 23:00 and 05:00, past the time of local flight activity, are assumed to be immigrants that were carried by winds and were transported to the trap location. Although rare, captures between 05:00 and 12:00 are interpreted as possible long-distant immigrants. This interpretation is then validated by cross-referencing with the location phenology (https://burps.budworm.ca) to determine if adults are expected in the area at a given date, and local temperature, as spruce budworm moths are unlikely to fly in cold temperatures (i.e., below 13°C for males and 17.5°C for females) (Sanders et al., 1978; Greenbank et al., 1980). Furthermore, capture numbers are also interpreted relative to recent day captures and activity across the approximately 30 traps in the network. Thus, high numbers of captures, even before 23:00, preceded and followed by much lower captures, are marked as a potential immigration event. Potential immigrants are then confirmed by looking at wind patterns during the time of capture, cross-referenced with areas where phenology indicates adults are present, and trajectory models are generated to evaluate potential sources of immigration (Stein et al., 2015; HYSPLIT, v.4.8- https://www.ready.noaa.gov/HYSPLIT.php). Furthermore, on average at a given location, SBW adult emergence occurs for a period of 2 weeks and local moth activity lasts 3 weeks, yet the influx of immigrants that remain active following dispersal can extend the duration of activity at any given site (Greenbank et al., 1980). Overlaps in immigration events and local emergence affect the reliability of captures around dusk, as surviving immigrants -which can live for about four days (Rhainds et al., 2022)- can be misidentified as locals without any additional tools for discrimination.

We selected three sites within the active defoliation area of the current BSW outbreak which have high population densities. These sites had the highest capture rates of the 2019 flight season: Forestville (48.738°N, −69.07783°W), St. Modeste (47.83683°N, −69.39133°W) and Baldwin (48.562°N, −66.1055°W) (Figure 1). The other sites were outside the current outbreak area and are mostly experiencing endemic (low-density) SBW population dynamics, but are predicted to be potential zones of future outbreak expansion. These sites had an overall low number of captures throughout the flight season: Zinc Mine Rd (50.273°N, −57.555°W), Arisaig (45.75°N, −62.16°W), and Inverness (46.196°N, −61.294°W) (Figure 1).

### Hydrogen isotope composition analysis

Before measuring δ²H values in SBW wings, we cleaned the wings in a 2:1v/v chloroform:methanol solution in three successive washes for 30 min, at least 3 h, and 15 min respectively, to remove glue (from the sticky roll), lipids, dust, and contaminants that could impact the *δ*²H values of the samples. The chloroform:methanol wash effectively removed these contaminants (Supplemental Figure 2). The wings were then air dried in a class-100 fume hood and stored in a glass vial. For each individual, we sampled 0.10 to 0.15 mg of wing tissues packed into silver capsules. The *δ*²H values of the non-exchangeable hydrogen of butterfly wings were determined at the Jan Veizer Stable Isotope Laboratory using the comparative analysis approach similar to Wassenaar and Hobson (2003). We performed hydrogen isotopic measurements on H_2_ gas derived from high-temperature (1400 °C) flash pyrolysis (TCEA, Thermo, Germany) of 0.15 ± 0.10 mg of wing subsamples, along with keratin standards: Caribou Hoof Standard (CBS; *δ*²H = −157.0 ± 0.9 ‰), Kudo Horn Standard (KHS; *δ*²H = −35.3 ± 1.1‰) (Soto et al., 2017), USGS42 hair (*δ*²H = −72.9 ± 2.2 ‰), USGS43 hair (*δ*²H = −44.4 ± 2.0 ‰) (Coplen and Qi, 2012), and two internal standards made of chitin material: ground and homogenized spongy moths (*Lymantria dispar dispar,* Linnaeus, 1758) (*δ*²H = −64 ± 0.8‰) and Alfa Aesar chitin (*δ*²H = −22 ± 1.2‰). The resultant separated H_2_ flowed to a Conflow IV (Thermo, Germany) interfaced to a Delta V Plus IRMS (Thermo, Germany) for *δ*²H analysis. The USGS42 hair sample was calibrated with a three-point calibration curve to the reference materials (i.e., CBS, KHS, and USGS43), while USGS42 and the two chitin internal standards were used as quality checks. The measured *δ*²H values for USGS42 (−72.5 ± 1.6‰, *n=4*), spongy moths (−−63.67 ± 0.52‰, *n=6*), and Alfa Aesar chitin (−19.25 ± 0.96‰, *n=4*) were within the reported value and uncertainty. The analytical precision of these measurements is based on the reproducibility of USGS42 and the chitin internal standards and is better than ± 2‰. All *δ*²H measurements are reported following the international scale VSMOW-SLAP.

### Strontium isotope ratio analysis

To remove the glue and other potential contaminants deposited on samples, we cleaned abdomen, thorax, and head tissue in a 2:1 chloroform:methanol solution in three successive washes for 30 min, 3 h, and 15 min respectively (Supplemental Figure 2). Samples were air dried in a class-100 fume hood and stored in glassine envelopes. Samples were then digested in concentrated nitric acid (16 M; distilled TraceMetal™ Grade; Fisher Chemical, Canada) for 15 minutes at 250°C using Microwave digestion (Organic High setting - Anton Paar Multiwave 7000, Austria). After digestion, the vials were limpid suggesting complete digestion. After drying the sample, 1 mL 6M HNO_3_ was added to each vial and then transferred to a 7 mL Savillex PTFE vial. An aliquot of 50 μL of the solution from each Savillex vial was pipetted to Labcon MetalFree_TM_ centrifuge tubes and diluted with 2 mL of 2% v/v HNO_3_. Sr concentration analysis was performed by Inductively Coupled Plasma Mass Spectrometry (ICP-MS) (Agilent 8800 triple quadrupole mass spectrometer) at the Department of Earth and Environmental Sciences, University of Ottawa. Calibration standards were prepared using single element certified standards purchased from SCP Science (Montreal, Canada).

The remaining ∼1 mL aliquot of the sample in the 7 mL Savillex PTFE vial was dried down and re-dissolved in 1 mL 6 M HNO_3_. The separation of Sr was processed in 100 μL microcolumns loaded with Sr-spec Resin_TM_ (100–150 μm; Eichrom Technologies, LLC). The matrix was rinsed out using 6 M HNO_3_. The Sr was collected with 0.05 M HNO_3_. After separation, the eluates were dried and re-dissolved in 200 μL 2% v/v HNO_3_ for ^87^Sr/^86^Sr analysis. The ^87^Sr/^86^Sr analysis was performed at the Pacific Centre for Isotopic and Geochemical Research using a Nu-Plasma II high-resolution multi-collector inductively coupled plasma mass spectrometer (MC-ICP-MS; Nu Instruments) coupled to a desolvating nebulizer (Aridus IITM, CETAC Technologies). The interference of ^87^Sr was corrected by subtracting the amount of ^87^Rb corresponding to the ^85^Rb signal. Instrumental mass fractionation was corrected by normalizing ^86^Sr/^88^Sr to 0.1194 using the exponential law. Strontium isotope compositions are reported as ^87^Sr/^86^Sr ratios. The reproducibility of the ^87^Sr/^86^Sr measurement for 5ppb NIST SRM987 is 0.71025 ± 0.00009 (1 SD, n = 138) and for 1.4ppb NIST SRM987 0.71019 ± 0.00011 (1 SD, n = 48). As Lepidoptera cuticle is made of chitin, we also used one chitin internal standard, 5ppb Alfa Aesar chitin 0.713959 ± 0.00009 (1SD, n=3).

### Statistical analyses

Since local individuals ought to have formed their tissues within the same location under a limited area, we expect putative local individuals captured at a given site to have similar *δ*^2^H values and ^87^Sr/^86^Sr ratios to each other. Furthermore, we expect isotope variation among these putative locals to have limited variation. At a given site, *δ*^2^H values usually fit a normal distribution (Hobson et al. 1999) whereas ^87^Sr/^86^Sr ratios can fit either a normal or lognormal distribution depending on the geological context (Bataille et al. 2020). Here, we assume a normal distribution for both isotopes at the site of collection. Unlike locals, immigrants arriving at a site may have heterogeneous origins - i.e., can originate in different sites across a broad spatial extent- and thus we expect them to have different isotopic values among themselves. Heterogeneous origins can also lead to non-normal distributions of values, and therefore we chose not to treat putative immigrants at a given capture site as if they were part of a homogeneous group. For these reasons, at each site and for each isotopic system, we performed Student’s t-tests on individual putative immigrants -instead of the group of immigrants as a whole- and compared them to the local population mean and standard deviation, as a way to determine whether they were true immigrants.

We further explored whether the isotopic composition of SBW can be used to discriminate between locals and immigrants using the combined information from dual hydrogen and strontium isotopes. Due to the uncertainty in the classification of observed individuals into local vs. immigrant groups, traditional methods for testing differences between groups (such as t-tests) have reduced power. While a t-test exists under uncertain group membership conditions (Bauer et al., 2021), this test relies on quantifying the group membership probability, and assumes equal variances and normality for both groups. We do not expect the variance of the local and immigrant groups to be the same, and since the immigrant group could have several different geographic origins, we cannot assume that the distributions of observed values for the immigrant groups are normal. With these violations of the typical assumptions required for comparing groups, along with the additional issue of uncertain group membership, we decided to examine robust estimates of location and scale for the assumed local groups in each location. By using a median to represent location and a scaled median absolute deviation (Rousseeuw and Croux, 1993) to represent scale, we anchor the estimates to central points in the group; ideally, these points are the most likely to actually belong to that group. We were unable to use robust estimates for the correlation between δ^2^H and ^87^Sr/^86^Sr, due to very small sample sizes of single observations with both measurements. We created bivariate normal ellipses to discriminate the local population signal from immigrant individuals. To this end, the median values for δ^2^H and ^87^Sr/^86^Sr were calculated across all putative local observations for each site, and these values were used as the location parameters. The median absolute deviations of these observations were also calculated, and scaled to use as estimates for sigma, the marginal standard deviations. To draw the ellipses, 100 points were sampled from bivariate normal distributions parameterized by these values along with the estimates of correlation, and the ggplot function stat_ellipse() was applied (package tidyverse Wickham et al., 2019). Only ellipses for putative local groups were generated since we do not assume that the values of our variables of interest in the immigrant groups are normally distributed. It is important to note that these plots are mainly for exploration and comparison, and are not used as a formal test or indication of group membership.

All analyses were performed using R 4.0.3 (R Core Team, 2020). We used the ggplot2 (Wickham, 2016) package for figures and the rgdal (Bivand et al., 2022) and raster (Hijmans, 2020) packages for spatial analyses.

## Results

### Trap capture over flight season

The frequency and number of captures in our automated trap network varied by site, in a manner consistent with expected population densities. Traps outside the SBW outbreak area had in general a lower number of captures, reflective of lower density populations (i.e., Arisaig and Inverness, but less so at Zinc Mine), whereas sites within the current outbreak zone had a high number of daily captures, as expected for high-density populations (i.e., Forestville, St. Modeste and Baldwin) (Figure 2A and 2B). Importantly, at Arisaig and Inverness (Nova Scotia), we sampled for putative locals early during the active season, and at a time point before evidence of immigration. At these two sites, immigration events on the 23^rd^ of July 2020 were confirmed by radar, all putative immigrants arrived before 23:00, and the event occurred about 10 days after any local activity had ceased at Arisaig, and at least two weeks after the time-point we sampled for locals. At Zinc Mine (Newfoundland - Figure 2C) we sampled putative locals from captures on the 30^th^ of July between 15:00 to 23:00, and captures from the 31^st^ of July between 07:00 to 15:00 and 15:00 to 23:00, whereas putative immigrants were sampled from a capture between 15:00 and 23:00 on August 3^rd,^ 2019, and confirmed as an immigration event by radar. Additionally, some mixing between putative locals and immigrants is likely to have occurred since on the night of July 30^th^ an immigration event, confirmed by light traps and radar, was detected at this site.

**Figure 2:**
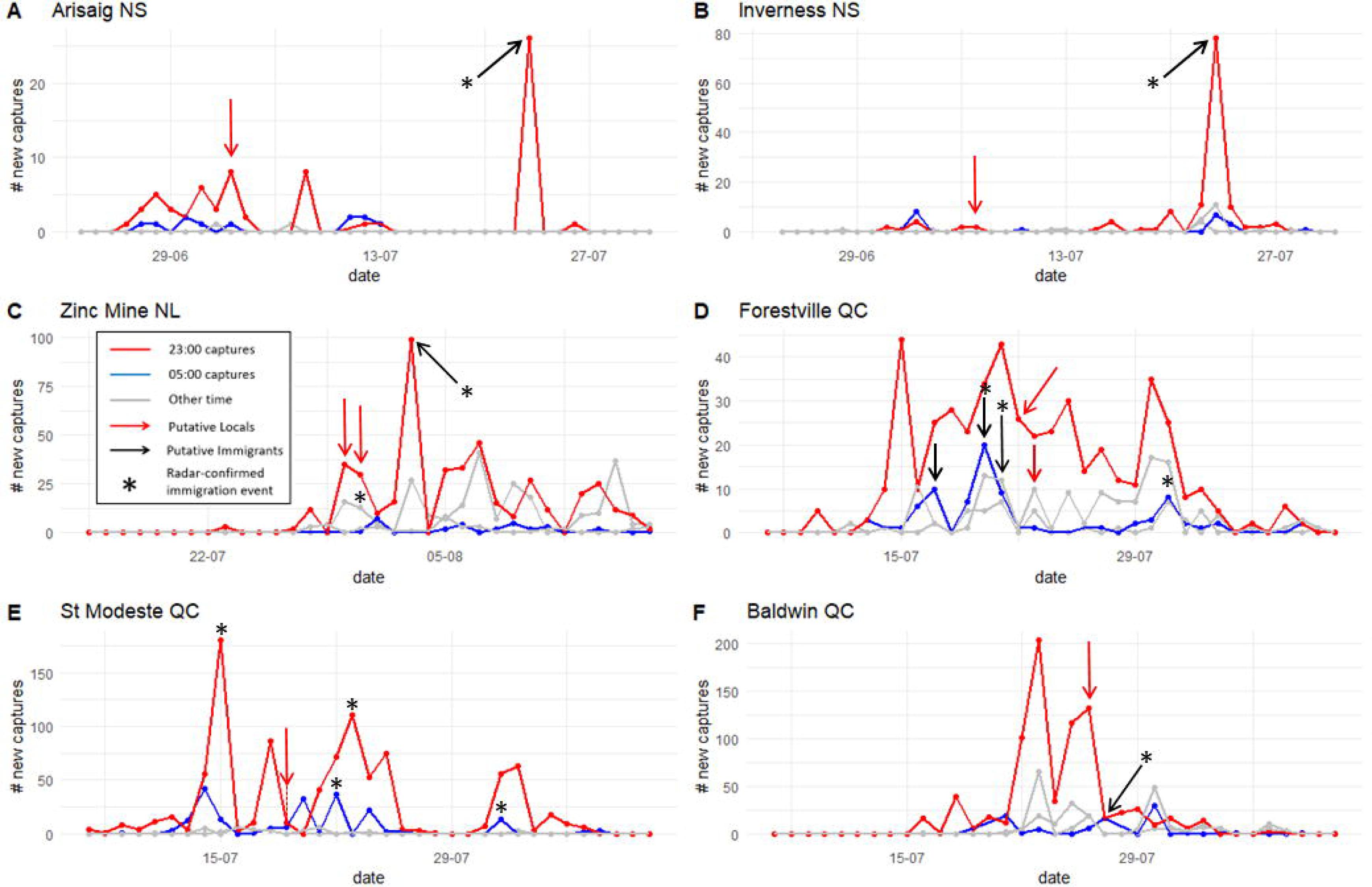
Number of eastern spruce budworm moths caught at each site on a given date and time. Each panel represents the flight season (June to August) captures of a given trap. Moths collected between 17:00 and 23:00 (red line) are often assumed to be putative locals, whereas moths captured between 23:00 and 05:00 (blue line) are interpreted as putative immigrants. Captures between 5:00 and 17:00 (grey lines) are more difficult to interpret and could be composed of late-arriving immigrants or early-flying locals. Arrows indicate time of sampling of local individuals (red arrows) and immigrant individuals (black arrows) for isotope analyses.

At St. Modeste and Baldwin (Quebec), putative immigrants were captured between 23:00 and 05:00 following dispersal events confirmed by radar (St. Modeste- 19^th^-20^th^ of July 2019, Baldwin-26^th^-27^th^ July 2019), and putative locals were sampled from captures between 17:00 and 23:00 the same night (i.e., July 19^th^ at St. Modeste and July 26^th^ at Baldwin) (Figure 2E and 2F). It is possible that dawn captures could be a mix of true local individuals flying close to dusk and early arriving immigrants. Additionally, although no radar-confirmed immigration events were detected previously at these two sites, high and sudden peaks on the 14^th^ and 15^th^ of July at St. Modeste, and the 23^rd^ of July at Baldwin, are suggestive of an earlier immigration event before our sample collection. At Forestville (Quebec - Figure 2D) we sampled putative immigrants from captures between 23:00 and 05:00 on the nights of the 16^th^ to the 17^th^ of July 2019 and 2 nights later on the 19^th^ to the 20^th^ of July. The peaks of July 17^th^ and 20^th^ at dawn are suggestive of an immigration event, and on July 20^th^ we also have radar confirmation of a mass flight reaching this area. We sampled putative locals from captures between 17:00 and 23:00 on the 22^nd^ of July 2019.

### Hydrogen isotope composition differences between local and immigrant SBW

The *δ*²H values of putative immigrant individuals were often different from that of the putative local group mean (Figure 3). In particular, in sites like Arisaig and Inverness, which are on the margins of the SBW distribution, and where putative locals were collected much earlier than any suspected immigration event, most putative immigrants were significantly different from and had lower *δ*²H values than putative locals (Figure 3A and 3B). At other sites, some individuals significantly diverged from the local population *δ*²H values and could be confirmed as immigrants (Figure 3C to 3F). In two instances, these immigrants had lower *δ*²H values than the average local population (Forestville and Baldwin - Figure 3D and 3F), whereas, in the other two, immigrants had diverging *δ*²H values relative to the putative locals (i.e., higher and lower values than the putative locals, Zinc Mine and St. Modeste - Figure 3C and 3E). At Zinc Mine, St. Modeste and Baldwin (Figure 3C, 3E and 3F) some putative immigrants were not significantly different from putative locals.

**Figure 3:**
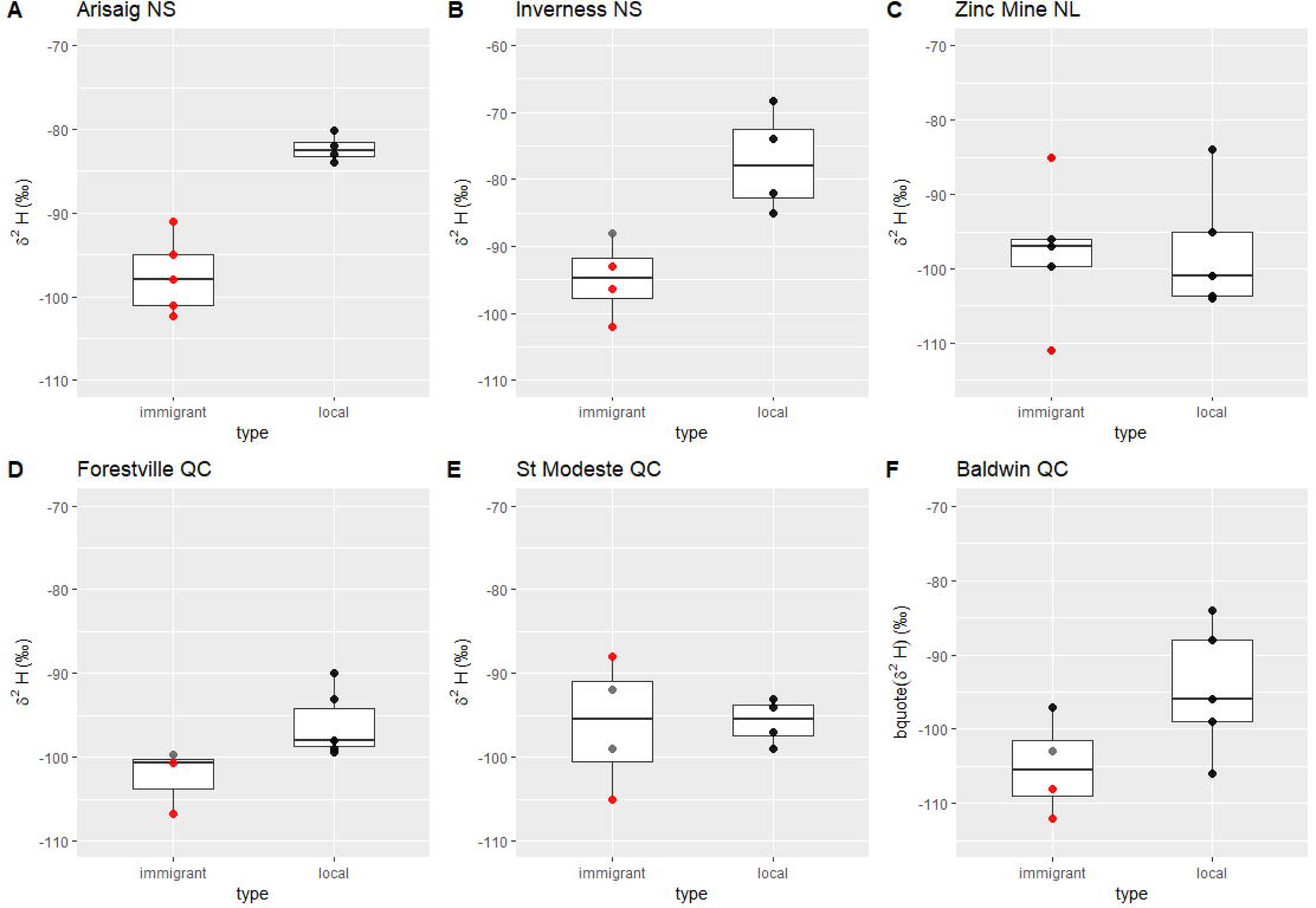
Box and whisker plot of *δ*^2^H values for immigrant and local SBW individuals by sampling site (population). Points represent individual samples. Putative local individuals are coloured black, immigrant individuals are coloured by whether they significantly differ from locals (red; t-test; p-value<0.05), were not significantly different (black; t-test; p-value>0.1) or were marginally nonsignificant (grey; t-test; 0.1>p-value>0.05).

### ^87^Sr/^86^Sr differences between local and immigrant SBW

When comparing individual immigrants to the mean local population value we found that ^87^Sr/^86^Sr ratios of putative immigrant individuals differed significantly from that of the putative local group mean in four of the six populations (Figure 4). As with δ²H values, Arisaig and Inverness showed putative immigrants having significantly different ^87^Sr/^86^Sr ratios than those of the putative locals. At Arisaig, immigrants developed in locations which have higher ^87^Sr/^86^Sr ratios (Figure 4A), whereas at Inverness immigrants arrived from sites with both higher and lower ^87^Sr/^86^Sr ratios (Figure 4B). Zinc Mine, on the other hand, did not show significant differences in the ^87^Sr/^86^Sr ratios of putative immigrant individuals and those of putative locals (Figure 4C). At St. Modeste and Baldwin, where putative immigrants were sampled immediately following the sampling of putative locals (23:00 to 05:00 and 17:00 to 23:00 respectively) the ^87^Sr/^86^Sr ratios of some putative immigrants differed significantly from that of putative locals, but several individuals showed no significant difference (Figure 4E and 4F). At Forestville, where immigration events had occurred five and two days before we sampled for local individuals, we found no difference between putative immigrant individuals and the putative local ^87^Sr/^86^Sr ratios, yet the range of putative local values is extreme in comparison with the isotope variation present in other putative local groups (from 0.71147 to 0.71440 - Figure 4D).

**Figure 4:**
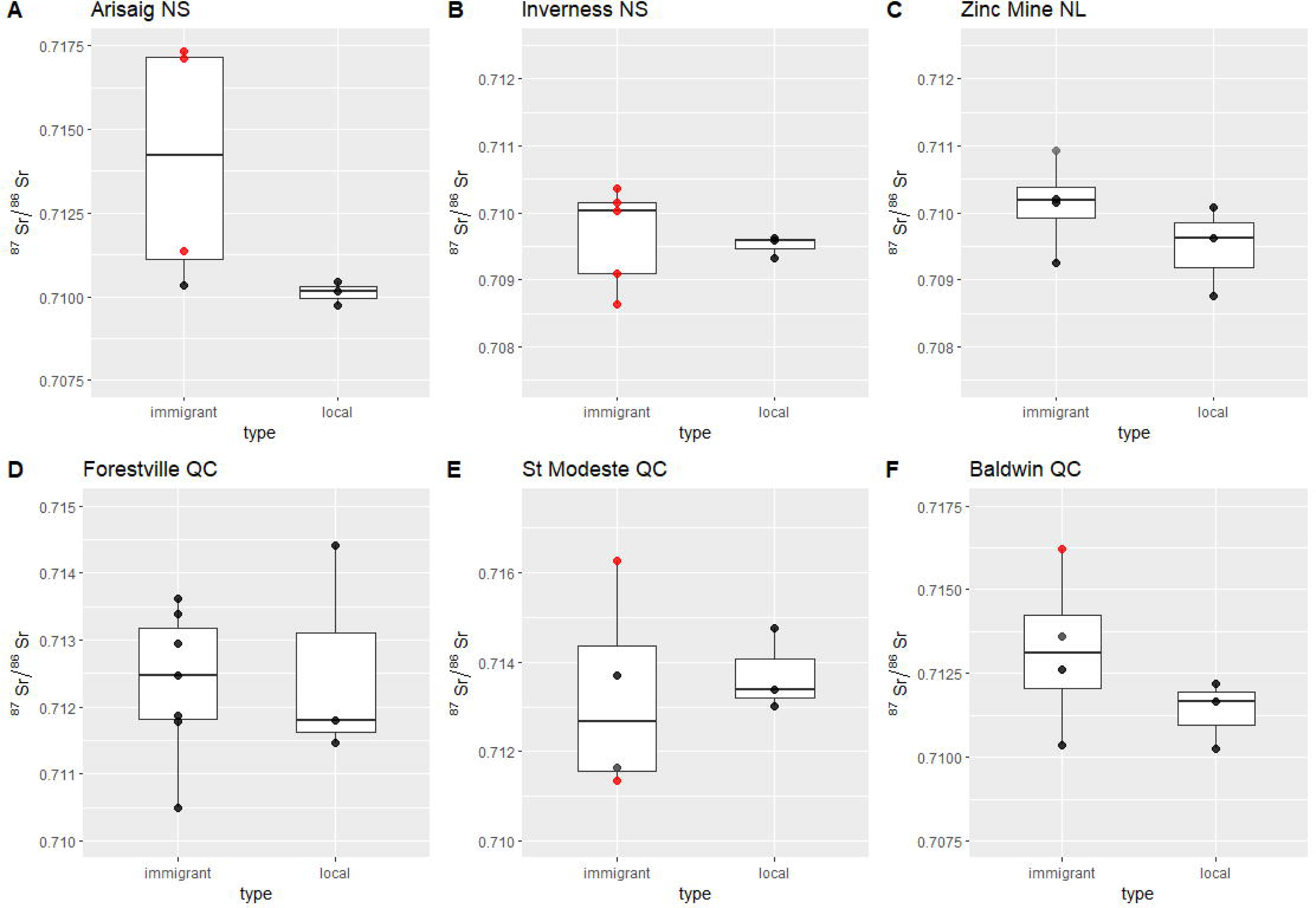
Box and whisker plot of ^87^Sr/^86^Sr ratios for immigrant and local SBW individuals by sampling site (population). Local individuals are coloured black, immigrant individuals are coloured by whether they significantly differ from locals (red; t-test; p-value<0.05), were not significantly different (black; t-test; p-value>0.1) or were marginally non-significant (grey; t-test; 0.105>p-value>0.05<=0.1).

### Dual hydrogen and strontium isotopes among local populations and immigrant individuals

Overall, in most instances, putative immigrants can be confirmed as true immigrant individuals since they do not fall in the same range of combined *δ*²H and ^87^Sr/^86^Sr values as locals (i.e., they fall out of the ellipses which are centred in the median value for both isotopes for putative locals -Figure 5). In particular, at Arisaig (Figure 5A) and Inverness (Figure 5B) all putative immigrants fall outside of the local confidence interval, showing that they developed at a site with different isotopic composition. Zinc Mine showed limited divergence among putative locals and immigrants, and our interpretations were limited by the scarcity of individuals we were able to sample for both *δ*²H values and ^87^Sr/^86^Sr ratios which did not allow us to create bivariate normal ellipses for this site. At St. Modeste (Figure 5E) and Baldwin (Figure 5F) some putative immigrants fall outside the ellipses whereas others are mixed with the putative local group and cannot be differentiated from putative locals, a result that is consistent with the single isotope data. The putative locals at Baldwin have a broad range of values for both *δ*²H values and ^87^Sr/^86^Sr ratios, a range of values unlikely for true locals. Forestville shows another interesting pattern, where several immigrants can be detected, but where putative locals also have more extreme values than immigrants (Figure 5D).

**Figure 5:**
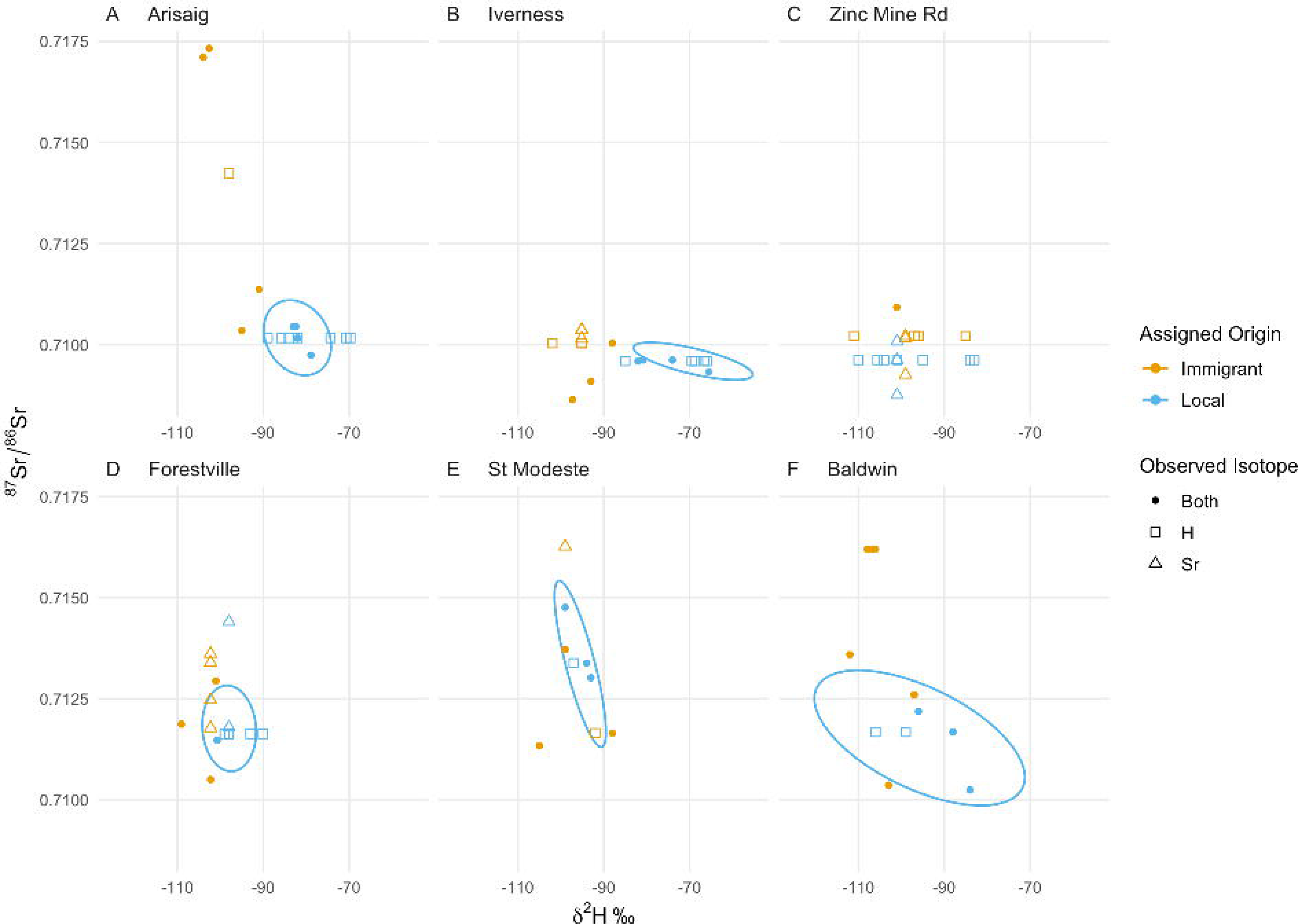
Dual isotope trait-space plot showing local population values (ellipse) and individual immigrants by site. Individuals with both a *δ*^2^H values and ^87^Sr/^86^Sr ratios are represented by filled symbols, individuals with only one isotope system measured are plotted at the median value of the isotope that was not measured. Points are segregated by putative immigrant/local status (*δ*^2^H-only values = squares, ^87^Sr/^86^Sr-only ratios = triangles). Ellipses are bivariate normal, centred in the media value of both isotope systems and used to visually discriminate between the local signal and immigrants. Zinc Mine only had one individual with dual isotope data, and we could not parametrize an ellipse.

## Discussion

The eastern spruce budworm (SBW) is the most severe pest of the boreal forest and causes millions of dollars in lost revenue to the Canadian economy during outbreak periods (MacLean, 2019). A key component of outbreak dynamics and spread is dispersal from areas of high SBW density (i.e., areas undergoing an outbreak) to areas of low density (i.e., undergoing endemic dynamics where SBW populations numbers are kept down through low larval survival rates while foliage searching and through the action of predators and parasitoids (Stedinger, 1984)). Spruce budworm moth dispersal has been challenging to monitor and manage, despite major improvements in the understanding of the ecology and pest control strategies of the species (Johns et al., 2019; MacLean, 2019). In this study, we use hydrogen and strontium isotopes to improve an approach currently used to determine migration events, which combines a network of automated traps that record the date and time of capture in discrete blocks, and evidence of local activity and dispersal from radar data, weather patterns and phenological records (see Methods).

We found that both *δ*^2^H values and ^87^Sr/^86^Sr ratios, individually and when combined, allow to discriminate between putative locals and immigrants. The isotope data generally supports the current classification approach based on time of capture (and other lines of evidence) particularly for identifying immigrants. Among all the putative immigrants analysed, at least one individual per site shows isotope data that differed significantly from that of putative local populations (Figures 3 and 4). Yet, in some instances, the flight behaviour and isotope methods do not support each other in identifying locals vs immigrants. In particular, when immigration events occurred before the collection of putative locals (e.g., up to approximately one week, e.g., Forestville, QC and Zinc Mine, NL) or when putative locals were collected immediately before an immigration event (e.g., Baldwin and St. Modeste, QC), the overlapping activity between locals and immigrants may lead to misclassification of trap captures (Figure 2). Our approach using combined *δ*^2^H values and ^87^Sr/^86^Sr ratios enhances the resolution of the above methods and allows to distinguish between SBW that completed their larval cycle at the site of capture (i.e., locals) vs. individuals from sites that differed in their isotopic composition (i.e., confirmed immigrants) (Figure 5). In particular, our novel isotopic tool allows to identify instances of potential misclassification and to make inferences about mixing of locals and immigrants in the population. Thus, *δ*^2^H values and ^87^Sr/^86^Sr ratios are an effective tool to confirm whether an immigration event has occurred at a given site and, with larger sample sizes, could also help determine the relative composition of locals and immigrants in a given site.

However, the confidence with which putative immigrants and locals were successfully identified based on capture-time and flight behaviour differed among sites, and reflected the nuances of local activity and immigration history at the sites of capture (Figure 2). Arisaig and Inverness (Nova Scotia) are located outside the range of the current SBW outbreak and have low, endemic, population density (i.e., comparatively low trap capture numbers) and experience less frequent immigration events (Figure 1). At these sites, only one immigration event was detected during the 2020 season and occurred several days (~10 days) after the peak activity of the local population (Figure 2A and 2B). The timing of the immigration event, close to dusk rather than after 23:00, suggests that these immigrants may have originated relatively close to (i.e., not hundreds of kilometres away) the capture sites, but the differences in *δ*^2^H values, in a few instances over 20‰, suggests that these individuals might have come from inland areas further north and away from the coast (Figure 3A and 3B). The ^87^Sr/^86^Sr ratios are generally higher and corroborate that the immigrants did originate in a site different from the capture (local) site. The higher ^87^Sr/^86^Sr ratios of immigrants at Arisaig (up to 0.717) support a northern origin on the Precambrian North American craton in Quebec, whereas the variable ^87^Sr/^86^Sr ratios of immigrants at Inverness suggest a more regional origin with ^87^Sr/^86^Sr ratios typical of the geologically younger Maritimes (Figure 3A and 3B). At both sites, dual isotope plots show clear discrimination among immigrant individuals and local values (Figure 5A and 5B). Furthermore, immigrants at both sites show a broad range of isotopic values suggesting that they do not come from a single area but instead have different geographical origins covering a likely broad spatial extent consistent with a large dispersal event. The remainder of the sites we sampled reveal a more complex pattern of immigrant and local distributions, highlighting some of the limitations of the flight-behaviour approach at high spatial resolutions, and underscoring the value of using isotopes to study SBW dispersal.

Using *δ*^2^H values and ^87^Sr/^86^Sr ratios showed that, at all sites, several putative immigrants were indeed immigrants, that is, these individuals had significantly distinct *δ*^2^H values and ^87^Sr/^86^Sr ratios from the putative local populations (Figures 3 and 5). This demonstrates that the combined approaches used to detect immigration (e.g., flight-behaviour and timing, radar, phenology, weather) are an effective tool to confirm whether an immigration event has occurred at a given area. Yet, our results also showed a few instances of putative local values falling well outside the range of expected local isotopic variation, and several instances of putative immigrants not being different from putative local values.

The first set of unusual outcomes points to likely misclassification of immigrants as putative locals. For example, the *δ*^2^H value of the putative local at Zinc Mine with a value of −84‰ (Figure 3C) and the putative local at Forestville with an ^87^Sr/^86^Sr ratio of 0.71440 (Figure 4D) place them as outliers and is well beyond the expectation of local variation that corresponds to a 20 km radius of variation in *δ*^2^H values from precipitation or in bioavailable ^87^Sr/^86^Sr ratios (Table 1A and 1B, Supplementary Table 3). One possible explanation for this possible misclassification is that following immigration to a given site, surviving immigrants then behave as locals on subsequent days, and lead to the mixing of immigrant and local individuals, an issue that is detected by our isotope tools but which current approaches, including population genetics (e.g. Lumley et al., 2020), have been unable to address.

Putative immigrant individuals that were not significantly isotopically distinct from local groups can be interpreted in several ways. First, they could be ‘true’ locals that were flying during the arrival of immigrants and thus are misclassified individuals. Although local flying activity and the likelihood of encountering the traps are expected to be highest around dusk (Greenbank et al., 1980), it is not known whether the influx of large numbers of immigrants and the likely spike in pheromones from arriving females, could influence local male activity and induce them to become active. Second, given that in our analyses immigrants are defined by comparison to putative locals, limited sample sizes of local individuals and misclassifications - which increase the estimate of the population variance - could decrease our ability to detect ‘true’ immigrants. For example, removing the outlier putative local from Forestville (i.e., the individual with an ^87^Sr/^86^Sr ratio of 0.71440) changes the putative local mean (± sd) and decreases the standard deviation from 0.71255 (±0.00161) to 0.71163 (±0.00023), making the difference between putative immigrants and locals more conspicuous. Increasing sample size and quality of local individuals (i.e., collections before the arrival of immigrants) will improve the precision of immigrant identification. Third, individuals that were not significantly different could still be immigrants, but which had originated at sites that have similar isotopic signatures to the local site or had travelled only a short distance. While both *δ*^2^H values and ^87^Sr/^86^Sr ratios are redundant independently (Bowen et al., 2005; Bataille et al., 2020), the redundancy decreases substantially when combining these isotope tools because of their independent patterns and scales of variations (Wunder, 2010). Globally, *δ*^2^H varies continuously at large spatial scales, with decreasing values as latitude increases and increasing distance away from coastal regions (Bowen and Revenaugh, 2003), and thus non-significant differences in *δ*^2^H values could be expected for short-distance migrants coming from climatically similar regions. ^87^Sr/^86^Sr ratios vary at a high spatial resolution, reflecting the age and composition of local geology, and thus even short-distance travellers could show significant divergence in their ^87^Sr/^86^Sr ratios. However, ^87^Sr/^86^Sr ratios are highly redundant at regional to global scale (Bataille et al., 2020). Therefore, similar dual hydrogen and strontium isotopes between locals and immigrants is unlikely.

We expected, based on their patterns and scales of variations, that local individuals would show limited variance in both *δ*^2^H values and especially ^87^Sr/^86^Sr ratios, since they originate in the same location and are expected to have limited dispersal (Sanders, 1983), relative to immigrant individuals that can have more spatially heterogeneous origins. We indeed find that, overall, ^87^Sr/^86^Sr ratios tended to show smaller variance among locals than immigrant individuals (in particular when we consider potential misclassification of putative locals - e.g. Forestville, QC) (Figure 4). With the exception of Arisaig, we did not detect this same trend in *δ*^2^H variance which is likely related to the low scale and continuous variations of this isotope. However, this observation of similar variance between immigrants and locals also underlines the relatively large *δ*^2^H variance of local individuals (SD=Arisaig: 1.66‰, Inverness: 7.6‰, Zinc Mine: 8.38‰, Forestville: 3.82‰, St. Modeste: 2.75‰, Baldwin: 8.76‰) well beyond the analytical uncertainty (<2‰). Typical sources of variation in local *δ*^2^H values, such as seasonal shifts, adult feeding and the formation of new tissue, are unlikely to be relevant to this species. SBW are short-lived and all individuals at any site have their life-history synchronised, with the emergence of adults - a reflection of developmental rates- occurring within a two-week period (Kucera, 1980; Sanders, 1985). Additionally, SBW does not feed as adults and we sampled wings for *δ*^2^H analysis, which have low metabolic activity (Lindroos et al. *this issue*), making seasonal variation in precipitation *δ*^2^H values unlikely to cause variation in the local moth values. Diet-induced variations at a given site are possible, as local SBW could feed on different trees with distinct water sources (e.g., precipitation, groundwater, lake, river water) depending on their rooting depth and position on the landscape. If the local landscape has multiple isotopically-distinct reservoirs, this could add further heterogeneity to the *δ*^2^H values that get integrated into spruce and balsam fir needles and then transferred to SBW, further increasing local *δ*^2^H value variance. Even within one single tree, there are several layers of needles grown at different years which might have *δ*^2^H differences. While SBWs are known to feed preferentially on fresh needles they can switch to older needles if they emerge too early in the spring to access young needles or when younger needles are not available (Régnière and Nealis, 2008). In any case, our data support the work of Hobson (1999; 2019) who found that the *δ*^2^H values of locally-raised monarch butterflies had a standard deviation of approximately 4.5‰ within any given site. Other studies that have analysed known-origin Lepidoptera also report large intra-site standard deviation sometimes >5‰ (e.g. Brattström et al., 2008; Satterfield et al., 2018). When considered together, our isotope results suggest that local moth individuals move further than the 100 m suggested by Sanders (1983). Isotope data rather suggest local mobility in the scale of kilometres but determining precisely the range and drivers of local SBW movements requires further investigation.

Our findings have fundamental and applied implications. First, we provide an independent validation that the combined use of automated traps, radar monitoring, phenology and weather data is effective at detecting immigration events in SBW, while underscoring the complementarity of using isotopes to validate and enhance the resolution of this trap network. Second, the difficulty in confirming immigration events in SBW means that current approaches to management rely heavily on widespread winter monitoring of larvae at sites with endemic population dynamics, to detect early immigration-driven population increases and to control population growth before it shifts to an outbreak stage (Early Intervention Strategy - Johns et al., 2019; MacLean, 2019). Although this approach is proving to be effective (MacLean et al., 2019), it is also expensive. The combined use of automated traps and isotopes provides a more targeted approach to confirm immigration events, estimate the relative proportion to which they contribute to the local adult population -a proxy for the following year’s reproductive output and population density- and allow cost-effective decision-making of where to invest in subsequent larval monitoring and eradication strategies. Finally, we confirmed that dual hydrogen-strontium isotopic tools can be applied for the geolocation of low-mass insect species with wing material for hydrogen isotope analysis below 150 *μ*g (~ two wings) and body mass for strontium isotopes below 3 mg. We also confirmed that combined *δ*^2^H values and ^87^Sr/^86^Sr ratios have enough discriminatory power to investigate the dispersal of these small insect species over small spatial scales (i.e., <100 km). To better understand dispersal dynamics in this system, our next step is to complete analyses on known-origin samples and develop SBW moth-calibrated *δ*^2^H and ^87^Sr/^86^Sr isoscapes for Eastern Canada, and then perform dual continuous isotope assignment of immigrant moths and reconstruct their origin and dispersal trajectory. The development of these isotope tools to study non-migratory wind-assisted dispersal behaviour of small winged insects will open new research avenues to investigate the mobility of many Boreal pest species (e.g., emerald ash borer, mountain pine beetle, spongy moth and Asian longhorned beetle). Dual isotope geolocation can help ascertain if the pest in question has been in a particular region for a long period (i.e., locally reared pest) or if it has recently arrived (i.e., long-distance migrants or invasives). It can also help identify their dispersal routes. Cumulatively, this information is key to develop more efficient early-eradication strategies (e.g., containment and spraying efforts) and to improve the cost-effectiveness and sustainability of forest management decisions by industry and government practitioners.

## Author Contributions

Conceived and designed the project: FD, JNC, CPB. Collected data: FD, JNC, KHP, MSR. Analyzed the data: FD, JNC, KS, CPB. Contributed reagents/materials/analysis tools: JNC, CPB. Wrote the first draft with input from coauthors: FD. Revised and edited the manuscript: FD, JNC, KS, KHP, MSR, CPB.

## Acknowledgments

We would like to thank our colleagues who helped collect samples at NRCan CFS [Emily Owens, Rob Johns, Gaetan Leclair and Jeff Fidgen] and in the provinces [Pierre Therrien and Jean-Jacques Bertrand (Quebec MFFP), Dan Lavigne and Troy Rideout (Newfoundland FIA)], Kerry Klassen and Paul Middlestead at the Jan Veizer laboratory for developing the low-mass hydrogen analyses, and Lihai Hu and Smita Mohanty for help with ICP-MS and strontium analyses.

## Supplementary Material

Supplementary Material should be uploaded separately on submission, if there are Supplementary Figures, please include the caption in the same file as the figure. Supplementary Material templates can be found in the Frontiers Word Templates file.

Please see the Supplementary Material section of the Author guidelines for details on the different file types accepted.

## Data Availability Statement

The datasets generated for this study and R code for analyses is archived with the Open Science Foundation [LINK].

