## Supplemental Materials for "Characterizing Eastern spruce budworm’s large-scale dispersal events through flight behavior and stable isotope analyses"

### *Supplementary Material*

#### 1 Supplementary Figure 1 – Example of trap captures and moth selection

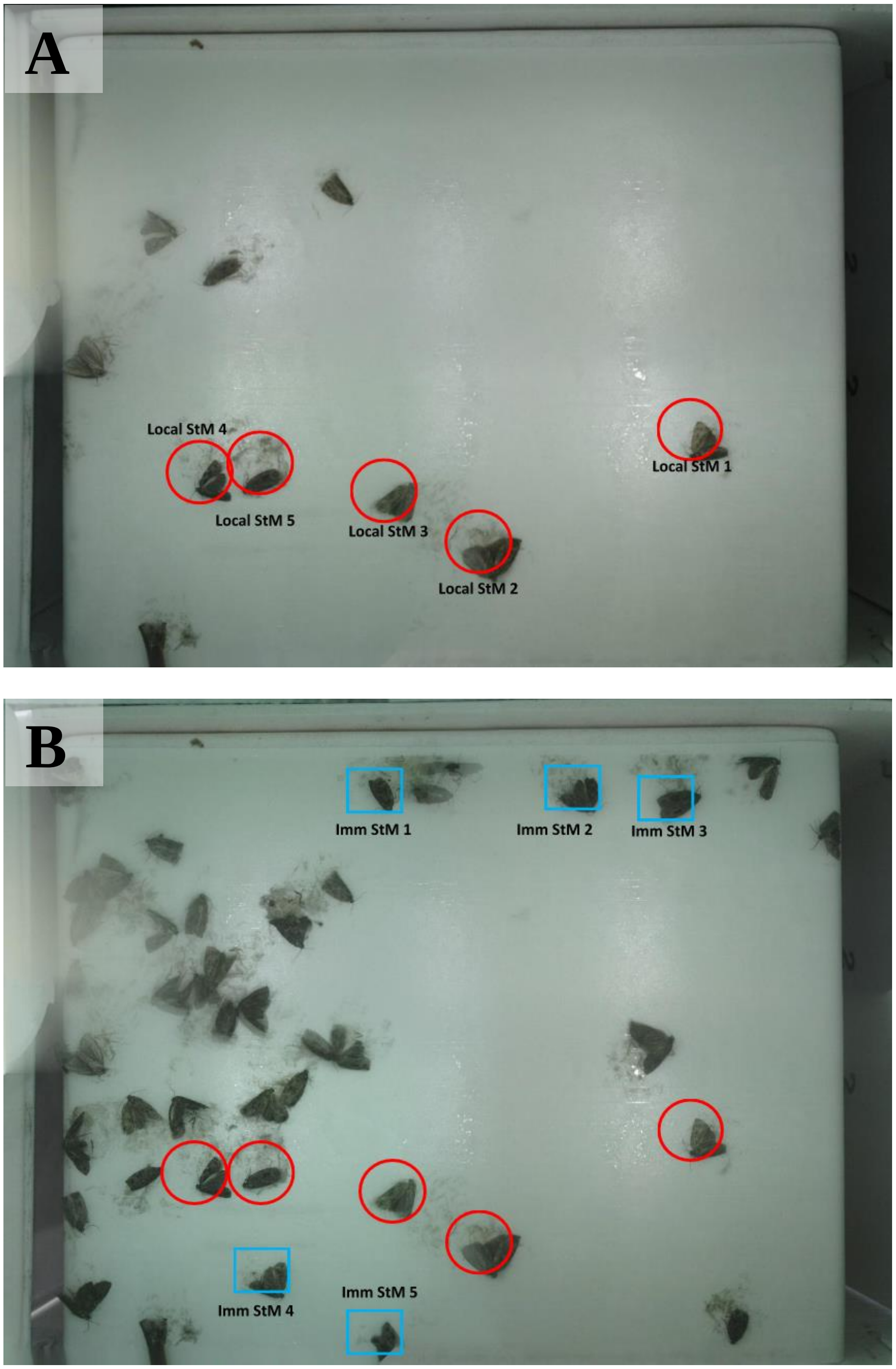


**Supplementary Figure 1.** (A) Trap image taken on the 19th of July 2019 at 23:00. The previous image, taken at 17:00, had no moths captured, after this a new segment of sticky paper was rolled.

The ten moths in the image, thus were captured between 17:00 and 23:00, the time associated with local individuals' flight. These individuals were assumed to be local and we sampled 5 of them (marked in red). (B) The following image was taken on the 20th of July 2019 at 05:00 and the trap now has 44 individuals, of which 34 arrived between 23:00 and 05:00, and are thus considered putative immigrants. Immigrants used for is+otopic analyses are marked in blue squares.

#### 2 Supplementary Figure 2 – Cleaning of moth tissues

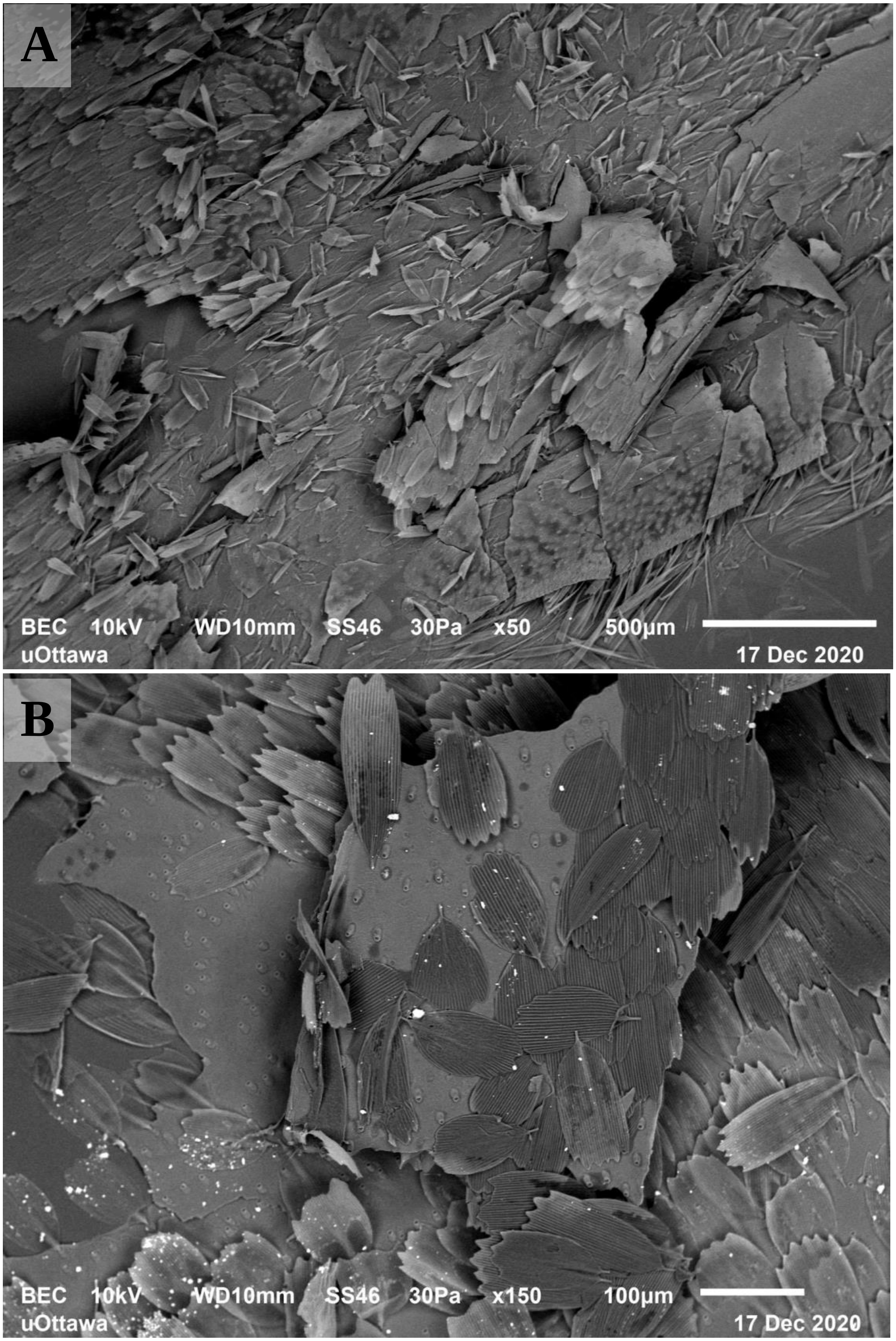

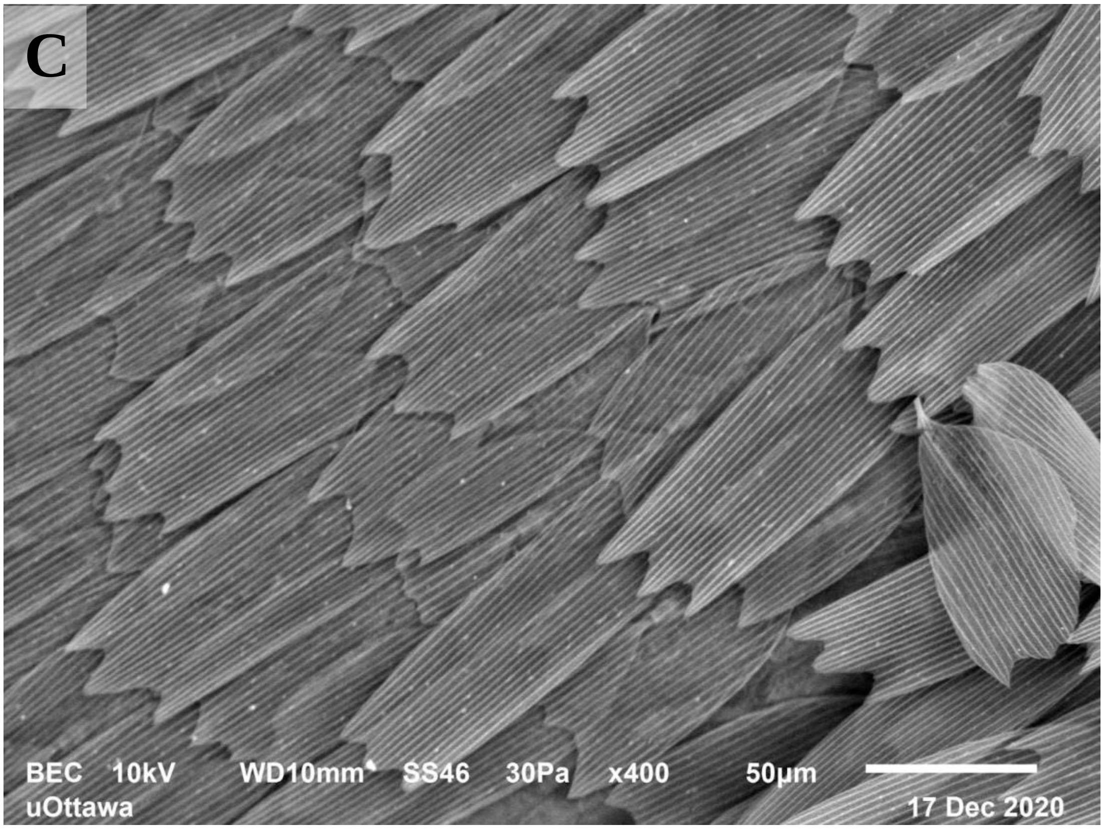


**Supplementary Figure 2.** Scanning Electron microscope images of moth wings (A) before and (B, C) after cleaning with 2:1 v/v chloroform:methanol in a cycle of three washes with a duration of 30 minutes, three hours, and 15 minutes. The solution was changed for each wash. Images show no glue residues or mineral particles attached to the wings after washing.

#### 3 Supplementary Table 3 – $\delta^2$H and $^{87}$Sr/$^{86}$Sr variation around trap sites

**Supplementary Table 3.** Variation in (A) $\delta^2$H values and (B) $^{87}$Sr/$^{86}$Sr ratios around automated capture traps. Columns report the average value, number of cells (n), and standard deviation (SD) of concentric buffer zones at 5, 10 and 20 km surrounding our automatic traps. $\delta^2$H values were extracted from the resampled 1 km$^2$ raster $\delta^2$H values precipitation isoscape from the assignR package v.2.1.1. and $^{87}$Sr/$^{86}$Sr ratios from a foliar isoscape for eastern Canada (Le Corre et al. *in prep.*). Both isoscape rasters had pixels with a 1 km resolution. We used package terra to create 1000, 5000 and 20000 pixel buffers around each trap site and proceeded to extract the value of each pixel. Note: The reported values are minimum values as they do not account for the uncertainty of the isoscapes at each pixel.

(A)

| Site | 5 km mean (n) | 5 km SD | 10 km mean (n) | 10 km SD | 20 km mean (n) | 20 km SD |
|---|---|---|---|---|---|---|
| Arisaig NS | -57.19 (76) | 0.35 | -56.88 (265) | 0.57 | -56.29 (857) | 0.85 |
| Inverness NS | -58.27 (79) | 0.41 | -58.57 (307) | 0.71 | -58.48 (967) | 0.69 |
| Forestville QC | -67.27 (78) | 0.41 | -67.57 (277) | 0.7 | -68.42 (883) | 1.34 |
| Zinc Mine NL | -64.71 (78) | 0.45 | -64.75 (262) | 0.61 | -64.78 (855) | 0.67 |
| Baldwin QC | -69.14 (79) | 0.19 | -68.86 (315) | 0.34 | -68.79 (1250) | 0.48 |
| St. Modeste | -64.56 (78) | 0.5 | -64.66 (312) | 0.9 | -64.92 (1114) | 1.23 |

(B)

| Site | 5 km mean (n) | 5 km SD | 10 km mean (n) | 10 km SD | 20 km mean (n) | 20 km SD |
|---|---|---|---|---|---|---|
| Arisaig NS | 0.712477 (46) | 0.000318 | 0.712203 (170) | 0.000559 | 0.711792 (646) | 0.000634 |
| Inverness NS | 0.711861 (66) | 0.000325 | 0.711759 (202) | 0.000333 | 0.711638 (737) | 0.000594 |
| Forestville QC | 0.715758 (32) | 0.000398 | 0.71609 (146) | 0.00037 | 0.716179 (645) | 0.000731 |
| Zinc Mine NL | 0.712764 (46) | 0.00037 | 0.712619 (139) | 0.000351 | 0.712537 (601) | 0.0004 |
| Baldwin QC | 0.717756 (78) | 0.001254 | 0.717796 (312) | 0.001369 | 0.717524 (1248) | 0.001855 |
| St. Modeste QC | 0.716989 (78) | 0.000728 | 0.717088 (309) | 0.000833 | 0.717366 (1006) | 0.000832 |
